# Visuomotor temporal adaptation is tuned to gamma brain oscillatory coherence

**DOI:** 10.1101/2020.06.29.177295

**Authors:** Clara Cámara, Cristina de la Malla, Josep Marco-Pallarés, Joan López-Moliner

## Abstract

Every time we use our smartphone, tablet, or other electronic devices we are exposed to temporal delays between our actions and the sensory feedback. We can compensate for such delays by adjusting our motor commands and doing so likely requires establishing new temporal mappings between motor areas and sensory predictions. However, little is known about the neural underpinnings that would support building new temporal correspondences between different brain areas. We here address the possibility that communication through coherence, which is thought to support neural interareal communication, lies behind the neural processes accounting for how humans cope with additional delays between motor and sensory areas. We recorded EEG activity while participants intercepted moving targets while seeing a cursor that followed their hand with a delay rather than their own hand. Participants adjusted their movements to the delayed visual feedback and intercepted the target with the cursor. The EEG data shows a significant increase in coherence of beta and gamma bands between visual and motor areas during the hand on-going movement towards interception. However, when looking at differences between participants depending on the level of adaptation, only the increase in gamma band correlated with the level of temporal adaptation. We are able to describe the time course of the coherence using coupled oscillators showing that the times at which high coherence is achieved are within useful ranges to solve the task. Altogether, these results evidence the functional relevance of brain coherence in a complex task where adapting to new delays is crucial.

**AUTHOR SUMMARY:** Humans are often exposed to delays between their actions and the incoming sensory feedback caused by actions. While there have been advances in the understanding of the conditions at which temporal adaptation can occur, little is known about the neural mechanisms enabling temporal adaptation. In the present study we measure brain activity (EEG) to investigate whether communication through coherence between motor and sensory areas might be responsible for one’s ability to cope with externally imposed delays in an interception task. We show evidence that neural coherence at gamma band between visual and motor areas is related to the degree of adaptation to temporal delays.

## INTRODUCTION

Humans are sensitive to temporal contingencies between actions and their sensory feedback [1]. For example, we easily notice a sudden increment of delay while operating a computer mouse because the new delay alters the expected temporal mapping between our hand movements and the predicted visual feedback. The prediction of sensory feedback based on motor commands is of paramount importance for forward models and seems essential to support accurate performance in the presence of inherent long neural delays [2, 3].

Predictions of sensory feedback become systematically inaccurate in the presence of perturbations of action feedback, be they spatial [4], temporal [5] or caused by a force-field [6]. Humans are able to adapt to new environmental conditions by adjusting their motor commands, and such adaptation likely requires establishing new predictions between the motor commands and the expected time of the corresponding sensory feedback [7]. Consequently, neurons between sensory (e.g. visual) and motor areas need to be coordinated.

Brain oscillations are thought to be an important component for the coordination of distant brain areas [8–10]. Among the different oscillatory components, several studies have proposed a key role of beta oscillations (13-25 Hz) in motor performance [11] and in sensorimotor corrections based on previous errors [12]. Beta power increases just before hand movement onset [13] and post-movement beta power has been postulated to reflect the uncertainty of estimations from internal models [14]. In relation to communication between different brain areas, beta band coherence involving multiple pathways (motor cortex, somatosensory areas and forearm muscles) has been reported in monkeys during a finger flexion task [15] with inputs from motor cortex (M1) to somatosensory areas (S1/PPC). This likely denotes the oscillatory efferent copy of the motor command from M1 setting an appropriate motor context to interpret sensory inflow. Synchronized oscillations influencing motor cortex from primary somatosensory and inferior posterior parietal cortices have also been shown in monkeys during motor maintenance behaviour [16]. On the other hand, low gamma (30-45 Hz) event related synchronization has been reported to start after onset of motor responses in humans, simultaneously to a desynchronization in the alpha band [17].

Time-sensitive communication between different areas that are relevant to solve a certain task (i.e. sensory and motor areas in the present study) relies on coherence. In fact, the coordination of distant brain regions is crucial in the performance of complex behaviours and might support key aspects in sensorimotor learning and adaptation [18]. To achieve this, any brain communication depending on oscillatory coherence between distant areas must cope with delays that could potentially undermine communication efficiency, affecting the phase difference (i.e. coherence) between them (see Figure 1A). In the case of visuomotor adaptation, some delays are inherent to the brain mechanisms underlying movement performance (such as neural integration or the axonal distance needed to integrate different brain areas), but others might arise from the environment (e.g. the delay between the computer’s mouse movement and the movement of the icon on the screen). Adaptation to imposed delays is possible [19] and has been shown to occur in both discrete actions like tapping [1] and continuous actions like tracking [20] or interception [21]. Importantly, in contrast to discrete actions, in continuous actions the motor command can be adjusted online by reacting to the delayed visual feedback of the on-going hand movement [5], which suggests an efficient and rapid coordination between motor and visual areas. In order to support this sensorimotor coordination and enable fast responses to additional delays, one must be able to rapidly process the temporal difference between the movement and the sensory consequences once the movement has started or after a correction. To achieve this, coherence between motor and sensory areas at high frequency bands might be relevant. Moreover, being able to adjust to imposed delays suggests that oscillatory motor activity at one time is phase-coupled with sensory oscillations at some time later. On a regular basis, it is possible that visuo-motor oscillations are phase-corrected for built-in delays (see Figure 1A). However, adapting to additional delays that are experimentally imposed, would require to correct and adjust the phases between two populations of neurons.

**Figure 1.**
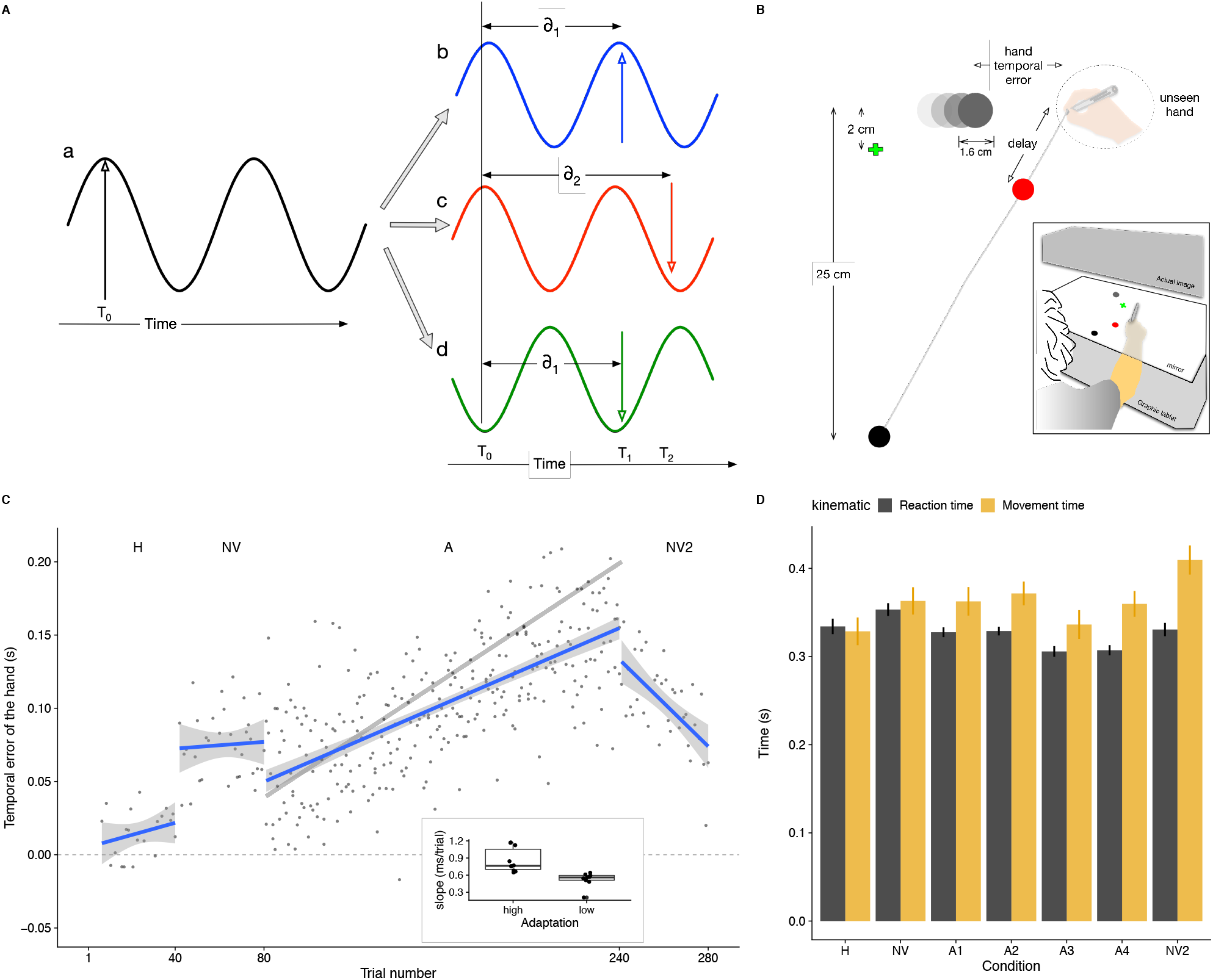
The role of phase coupling in temporal adaptation, experimental setup, and behavioural data. (A) Adapting to new visuomotor delays requires correcting for phase differences between motor and sensory areas. Suppose oscillatory activity in motor neurons (a, black oscillations) and sensory neurons (b, c and d). Efficient communication through coherence (i.e. small phase difference in the oscillations) means that the respective phases are corrected for neural delays (δ) and that information sent by ‘a’ is received by ‘b’, ‘c’ and ‘d’ when their oscillations are at a maximum. In the example, the information is sent from ‘a’ at time T_0_ (black vertical line) and arrives after a delay δ_1_ to ‘b’ when ‘b’ has an oscillation peak (vertical red line at time T_1_). Any additional new delay (δ_2-_ δ_1_) imposed by means of a perturbation might not be accounted for and communication between neurons ‘a’ and ‘c’ would not be efficient despite them being in phase (black and red oscillations). Thus, this would have the same result as if the neurons were not in phase (a and d: black and green oscillations). Therefore, an additional delay would introduce a corresponding phase difference. Adaptation would then correct for this phase difference and correct for the additional delay. (B) Participants had to intercept horizontally moving targets by sliding a stylus they held on their hand over a graphic tablet while fixating on a green cross. The experiment was divided in four conditions differing on the visual feedback of the ongoing movement. If participants used the visual information from the cursor in the A condition, they would cross the target’s path with the unseen hand ahead of the target in order to see the (delayed) cursor hitting the target. The inset represents the setup schematically. (C) Mean temporal error across participants as a function of trial. The grey line in condition A denotes the expected pattern of temporal errors for full adaptation. The blue lines are linear fits to the data points, and the grey areas show the 95% confidence interval for each condition. The inset shows the degree of adaptation(the individual slope in the A condition) of each participant segregated in high and low adaptors. (D) Mean reaction and movement time (black and yellow columns, respectively) across participants for the different conditions (with A divided in 4 blocks). Error bars correspond to the 95% confidence intervals within participants for each condition.

To understand how the brain copes with built-in neural delays while keeping high levels of coherence, theoretical models have used zero-lag long range synchronization approaches [22, 23]. Recent studies have also proposed approximations to correct for these delays based on coupled oscillators, being the Kuramoto oscillator [24, 25] a well-known coupling oscillator model. Here we use a spatial-embedded Kuramoto oscillator [25], which approximates neural delays, to describe how neural oscillators can give rise to coherence levels in time intervals that could support rapid communication in the context of an interception task with additional delays.

Notwithstanding the importance of this issue, there are no experimental studies relating the level of coherence between motor and sensory areas and temporal adaptation in a goal directed task. To this point, adaptation to imposed delays between actions and incoming feedback provides a touchstone for the role of oscillatory coherence and the feasibility of coupling mechanisms for correcting additional delays on top of the ones built-in in the nervous system. The goal of the present study is to determine whether the brain oscillatory coherence between visual and motor areas is functionally related to the human kinematic behaviour when adapting to delays, and if the time course of phase coupling in Kuramoto oscillators would allow these oscillators to be a feasible mechanism to account for the same time course of experimental brain coherence. With this aim, we studied brain oscillatory activity through EEG recorded during an interception task. The task consisted of different conditions that differed in the visual feedback provided of the on-going hand movement: participants could either see their hand while moving, see a delayed cursor representing their hand, or have no feedback of their hand during its movement. We expect to see higher levels of coherence in high frequency bands for higher levels of adaptation, and to describe the time course of the coherence using coupled oscillators.

## MATERIALS AND METHODS

### Participants

A total of 20 naive right-handed participants (14 females; range of age between 18 and 30) took part in the experiment. All participants had normal or corrected-to-normal vision and none had evident motor abnormalities. All participants gave written informed consent in advance and received a monetary compensation (30€) for their participation. The study was approved by the local research ethics committee.

### Experimental design and statistical analyses

#### Stimulus and procedure

Participants sat in front of a digitizing tablet (Calcomp DrawingTablet III 24240) that recorded the movements of a hand-held stylus while performing an interception task (Figure 1B). Stimuli were projected from above with an inverted LED LCD (ASUS) monitor positioned 45 cm above the tablet surface (inset in Figure 1B). Images were projected at a frame rate of 90 Hz with a resolution of 1280 × 960 pixels. A half-silvered mirror was positioned halfway (22 cm) between the monitor and the tablet, such that stimuli appeared to be in the same plane as the tablet. LEDs under the mirror allowed us to control the visibility of the hand: if the lights were on the hand was visible, but if the lights were off the hand was not seen. A custom program written in C and based on OpenGL controlled the presentation of the stimuli and registered the position of the stylus at 125 Hz. The setup was calibrated by aligning the stylus with the position of five dots appearing consecutively at the center and the corners of the screen, at the beginning of each session.

Participants were instructed to intercept a horizontally moving target by sliding the hand-held stylus across the drawing tablet. The target was a 16 mm-diameter white disk that moved on a black background (shown grey on white in Figure 1B). To start each trial, participants had to move the stylus to a starting position indicated by a 6 mm-diameter white dot (black disk at the bottom of Figure 1B) that was located 25 cm closer to the participants’ body than the target’s path. As soon as the hand was at the starting position, a green 3 mm-length cross indicating where participants had to fixate their gaze during the trial appeared at the vertical midline and 2 cm below the target’s path, and an auditory cue indicated that the trial was about to start. One second after the sound, a target appeared 10 cm either to the left or to the right of the vertical midline and moved horizontally in the opposite direction (i.e. if the target had appeared on the left it moved rightwards, and vice versa). The target moved at a constant velocity of either 20, 27, or 35 cm/s. The velocity and direction of the target were randomly chosen on a trial-by-trial basis.

The experiment consisted of 280 consecutive trials in which the visual feedback of the hand changed in four different ways (different experimental conditions). In the first 40 trials, participants saw their hand while moving (Hand condition - H). For the next 40 trials, the lights beneath the mirror were off and visual feedback of the hand was not provided (No vision condition - NV). The adaptation condition (A condition, 160 trials) followed the NV condition. In this condition, participants did not see their hand but saw a cursor that consisted of a 6 mm-diameter red disk that followed the movement of their hand. The cursor was delayed with respect to the unseen hand, and such delay was increased by 1 ms per trial [5]. The minimum constant delay of the system was 40 ms, so the additional delays were added on top of such minimum (thus, the cursor was delayed by 200 ms with respect to the unseen hand in the last trial of the A condition). The use of an adaptation paradigm [5] prevented participants from being aware of the manipulation and adopt strategic processes [26, 27]. During the last 40 trials of the experiment feedback of the ongoing movement was removed again (No vision condition - NV2). In order to help participants to find the starting position at the beginning of each trial in the NV and NV2 conditions, a cursor representing their hand was shown only when they moved in the space below the starting position.

#### Electrophysiological recording

EEG was recorded using 29 tin electrodes mounted in an elastic cap in standard positions (BrainAmps (c) -- Fp1/2, Fz, F7/8, F3/4, Fc5/6, Fc1/2, Cz, C3/4, T3/4, Cp5/6, Cp1/2, Pz, P3/4, T5/6, Po1/2, O1/2). Vertical eye movements were monitored with an electrode at the infraorbital ridge. Electrode impedances were kept below 5 kΩ during all the experiment. The electrophysiological signals were recorded at a sampling rate of 250 Hz with an online bandpass filter 0.01 to 70 Hz and were referenced to the lateral outer canthus of the right eye. The computer used to display the stimuli and the EEG recording unit were synchronized via a DAQ labjack U12 (Labjack (c). Lakewood, CO, USA) that was used to trigger signals to identify the stimulus onset and offset of each trial.

#### Behavioural data analyses

To analyse the hand movements, the data of the stylus’ position that participants held in their hand while moving was digitally low-pass filtered with a Butterworth filter (applied in both directions to avoid a phase shift), with a cut-off frequency of 10 Hz. Further analyses were based on this filtered data. The tangential velocity was computed by differentiating the smoothed positions. For each trial, we computed offline the hand and the cursor reaction time (RT) by using a velocity threshold of 4 cm/s. We also calculated when the hand and the cursor crossed the target’s path. The movement time (MT) was defined as the difference between when the hand started moving and when the hand (or the cursor in the A condition) crossed the target’s path. We analysed the effects of the different feedback conditions on these variables by fitting linear mixed-effect models (LMMs), which allow to easily incorporate trial effects in the same analysis because they do not require prior averaging. To do so, we used the lmer function (v.1.0–6) [28] from the R software. Linear contrasts were performed on the LMMs to conduct pairwise comparisons between conditions based on z scores with the glht function in R [29]. When reported, the p values obtained in the pairwise comparisons were adjusted using the Hochberg method [30].

In order to see whether participants relied on visual feedback of the cursor to try to intercept the target we looked at the temporal errors between the hand and the target at the moment the hand crossed the target’s path [5, 21, 31]. The temporal error then is defined as the time difference between when the center of the target and when the hand cross the point at which the hand crosses the target’s path (Figure 1B). A temporal error of zero means crossing the target through its center with the hand. Positive values would indicate that the hand crossed the target’s path ahead of the target, and negative values that the hand crossed the target’s path behind the target. When a delayed cursor is shown (A condition, Figure 1B) the temporal error is used as a measure of the degree of adaptation to the imposed delay. If there is a delay between the hand and the cursor and one wants to see the cursor hitting the moving target, the hand will have to cross the target’s path ahead of the target. If the temporal error between the hand and the target is equal to the imposed delay between the hand and the cursor, the cursor will cross the target through its centre. Since the delay between the target and the cursor was increased by 1 ms/trial, adapting to the delay would imply hitting further ahead of the target in each trial, with a slope of 1 ms/trial in case of full adaptation (gray line in Figure 1C).

#### EEG analyses

The EEG signal was analyzed offline with the eeglab package [32] and customized scripts in MATLAB. The EEG signal was re-referenced off-line to the mean of the activity at the two mastoid processes and was filtered offline using a bandpass filter from 0.1 Hz to 45 Hz. Single trials were time-locked to the hand movement onset and 3.5 seconds epochs, from −2 s to 1.5 s, were created. The period between 700 and 500 ms before the hand movement onset was taken as a baseline. In order to remove unwanted additional signals, such as oculomotor artifacts and facial muscle activity, Independent Component Analysis (ICA) was computed as an artifact reduction approach [33] for each participant. This transformation generated independent source components (ICs) that isolated artifacts that we manually removed from the neural activity components, reconstructing a cleaner EEG signal. After reconstructing the signal, we had to exclude two participants (one female) from the EEG analyses because the cleaned signal was not good enough for further analyses. After excluding these participants, we also checked for large shifts in voltage (which could also be due to oculomotor artifacts) within each epoch. Trials that presented an amplitude of more than ± 100 μV were automatically rejected off-line (from the remaining 18 participants, less than 4% of the EEG epochs were rejected). At this preprocessing stage, both sets of data (EEG and behavioural) were carefully matched trial by trial, which allowed to later segregate the different experimental conditions. In order to convert the EEG time series data into its time-frequency domain, the last step was to apply the complex Morlet wavelet at each trial (5 cycles, frequencies from 1 to 40 Hz, linear increase). The phase angle for each time and frequency was computed for a time window from −900 to 1000 ms relative to the hand movement onset. Then, to compute the phase-based coherence between pairs of electrodes, we used the Intersite Phase Clustering index (ISPC [34] also known as Phase-Locking Value [35]), which is defined as the length of the average vector of a distribution of phase angle differences over trials of the same experimental condition at each time-frequency point:

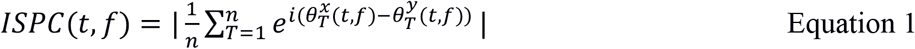

where *n* is the number of trials, and 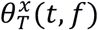 and 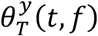 are phase angles of the complex wavelet coefficients from electrodes *x* and *y* at trial *T* for a specific time point *t* and frequency *f*. Due to the digitalization of the EEG signal, each time step corresponds to 4 ms. The ISPC resulting value ranges from 0 (for completely asynchrony) to 1 (for perfect connectivity). The frequency range was divided in different frequency bands for the analysis and statistic interpretations: 4-8 Hz (theta band), 9-15 Hz (alpha band), 16-25 Hz (beta band), and 26-40 Hz (gamma band).

To avoid background amplitude activity acting as a distraction, we selected a baseline between 700 and 500 ms before the hand movement onset and the percentage of ISPC change respect to this baseline was obtained per participant and condition. Motivated by our hypothesis, we restricted the ISPC respect to the baseline analysis to the activity recorded at the pair of electrodes located at near motor (C3/4) and visual (O1/2) areas. In other words, ISPC was computed only for the pairs of electrodes: C3 and O1, C3 and O2, C4 and O1, and C4 and O2. Since participants were fixating in a cross located at the sagittal midline, and target appearance was randomized in order to have 50% of the targets appearing at the left side of such midline and moved rightwards (the other 50% appeared at the right side of the sagittal midline and moved leftwards); we assume symmetry on the relevant visual information acquired by both electrodes located in the visual cortex (O1/2). For this reason, we averaged both pairs of ISPC respect to the baseline for each motor hemisphere. Hereafter, we refer to this averaged ISPC with respect to the baseline between C3 and O1 and C3 and O2, as the coherence between C3 and O1/2 (the same for the coherence between C4 and O1/2), for simplicity.

## SIMULATIONS

### Kuramoto oscillator

We simulated the degree of phase coupling between neurons using a Kuramoto model. We simulated a fully connected network of oscillators in which half of the oscillators (group 1) were fully bidirectionally connected with the other half (group 2) with the corresponding delay in analogy with motor neurons being connected with distant sensory neurons. For oscillators within the same group the delay was set to 0.

In the Kuramoto model the time evolution of the phase *ϕ* for *N* oscillators are given by the following equation:

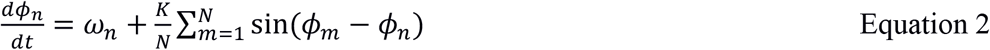

The first term (ω_*n*_) denotes the frequency of the oscillator *n* and it would correspond to the frequency with which a neuron fires spontaneously. The second term denotes how the oscillators interact. The constant *K* determines the strength of the coupling between oscillators *n* and *m* and, therefore, the tendency to be synchronized. However, Equation 2 does not consider the neural delays, that is the time that it takes for the information to travel from oscillator *m* to *n* (or vice versa). As shown in Figure 1A, it is important to consider this delay because it could introduce a phase shift. The dynamics of coupled Kuramoto oscillators with transmission delays can be approximated by the following expression [25]:

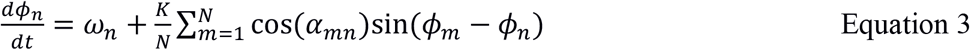

where *α*_*mn*_translates the time delay *τ*_*mn*_between oscillators *m* and *n* into a corresponding phase offset. In order to intuitively grasp the rationale of equation (3), suppose that the phases of two oscillators *m* and *n* are pi/2 and 0 respectively. This would translate to −1 in the term sin(*ϕ*_*m*_ − *ϕ*_*n*_). However, suppose that there is time delay between the two oscillators whose corresponding phase *α*_*mn*_ matches exactly the phase difference *ϕ*_*m*_ − *ϕ*_*n*_. In this example, *α*_*mn*_ = *pi*/2 and the time delay would be 6.25 ms. When multiplying the interaction term with cos(*α*_*mn*_) the result is very close to zero denoting that no phase correction is needed, since the delay between the two oscillators already compensates the phase difference.

We used equation (3) on our simulations with different number of oscillators, frequencies and delays. In different simulations, the number of oscillators could be *N* = {2, 10, 100, 1000}, and the frequency of the *N* oscillators was set to ω ={6, 12, 25, 40} Hz. These values were crossed with the delays: *τ* = {0, 50, 100, 150} ms. The combination of these variables resulted then in 5×4×4 (80) different simulated combinations. We used a single value of K (K=0.1, partial coupling) in all simulations and each combination was simulated 1000 times. We used the classical Runge-Kutta 4th Order Integration algorithm (RK4) to solve Equation 3. The initial phase *ϕ*_*m*_ of any oscillator *m* was set at random in the interval [−*π*, *π*] from a uniform distribution and *α* was the phase from the corresponding simulated time delay *τ* and oscillation frequency *ω*:

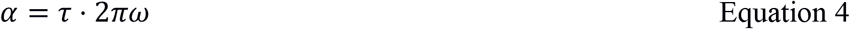

At each time step, the distribution of the of phases *ϕ*_*m*_ denotes the degree of coherence which would be minimum when the phases are uniformly distributed corresponding to a circular variance of 1. In order to obtain a measure of the coherence we computed the circular variance *V* (ranging from 0 to 1) and the measure of coherence (*R*) or synchronization was *R* = 1 − *V*. Since we are basically interested in the time course of R, this value was computed at each time once a whole solution for the step (up to 350 ms).

## RESULTS

### Behavioural

The temporal errors participants made in trying to hit the target differed depending on the feedback condition (Figure 1C). In the H condition, the mean temporal error across trials was 15 ms (*t* = 4.17, p = 0.0005), but this error did not significantly increase across trials (slope = 0.4 ms/trial, *t* = 1.2, p = 0.24). Considering that the target’s radius was 8 mm, missing the target by 20 ms would translate into a spatial error of 7 mm for the highest target’s velocity (35 cm/s) meaning that participants, on average, did not miss the target in the H condition.

In agreement with previous studies [5,21], removing the visual feedback (NV condition) made participants cross the target’s path ahead of the target (mean of 68 ms, t = 3.1, p = 0.004) but as in the H condition this error did not change across trials (0.1 ms/trial, t = 0.3, p = 0.75).

Unlike in the previous conditions, in the A condition the temporal error did increase significantly across trials (0.68 ms/trial, t = 18, p < 0.0001). The imposed delay increased by 1 ms/trial (grey line in Figure 1C condition A), so if participants aimed at seeing the cursor cross the target, they had to cross the target’s path ahead of the target. Even though the pattern of temporal errors denote adaptation to the delay, the difference between the blue and the grey line indicating what errors would make the cursor cross the target indicates that participants did not fully account for the imposed delay. Previous studies reporting a lack of complete adaptation have proposed forgetting mechanisms [37] or increased measurement noise [38] as possible causes to account for this incomplete learning. In our case, the lack of full adaptation may have also been caused by the use of the fixation cross to stabilize the gaze, which could have influenced the precision with which participants guided the cursor [39]. In order to check how movements were adjusted to compensate for the delay we looked at the reaction and movement times. One possible way to compensate for the temporal difference between the hand and the cursor is by starting to move earlier and then having more time to move and adjust the ongoing movement [5]. To evaluate changes in performance during adaptation, the 160 trials of the A condition were divided into four blocks (A1 to A4) of consecutive 40 trials so that all blocks/conditions had 40 trials per participant. Figure 1D shows that indeed the reaction times (black columns) became shorter (difference between A4 and H condition of 27 ms, z = 6, corrected p < 0.0001) and movement times (yellow columns) became longer (difference between A4 and H condition of 29 ms, z = −3, corrected p = 0.029) with the increasing delay.

During the last 40 trials visual feedback of the ongoing movement was removed again (NV2) and the temporal error decreased at a rate of 1.4 ms/trial (t = −4.6, p < 0.0001) until reaching a similar value (70 ms) to the error in the NV condition (Figure 1C). Returning to error values similar to the ones obtained before being exposed to the delayed cursor indicates that errors in the A condition were due to the use of the delayed cursor to guide one’s movements.

### Sensorimotor phase-base coherence

Figures 2 and 3 show the averaged coherence between C3 (contralateral motor response) and O1/2 and between C4 (ipsilateral motor response) and O1/2, respectively. In both figures the average is across participants. Each panel corresponds to a different condition (with the A condition divided in 4 consecutive blocks, each having an average delay increment of 40 ms with respect to the previous panel). The time value of zero (black-solid vertical line) corresponds to the hand’s RT. Note that all trials were aligned at zero for the hand RT before analyses. The different vertical lines correspond to the average across trials and participants of (from left to right) the stimulus onset (dotted line), the hand’s RT (continuous line at time 0), the hand’s MT (dashed line) and the cursor’s MT (red line). As mentioned in the methods section, the frequency values selected to define each band were 4-8 Hz for theta, 9-15 Hz for alpha, 16-25 Hz for beta and 26-40 Hz for gamma.

**Figure 2.**
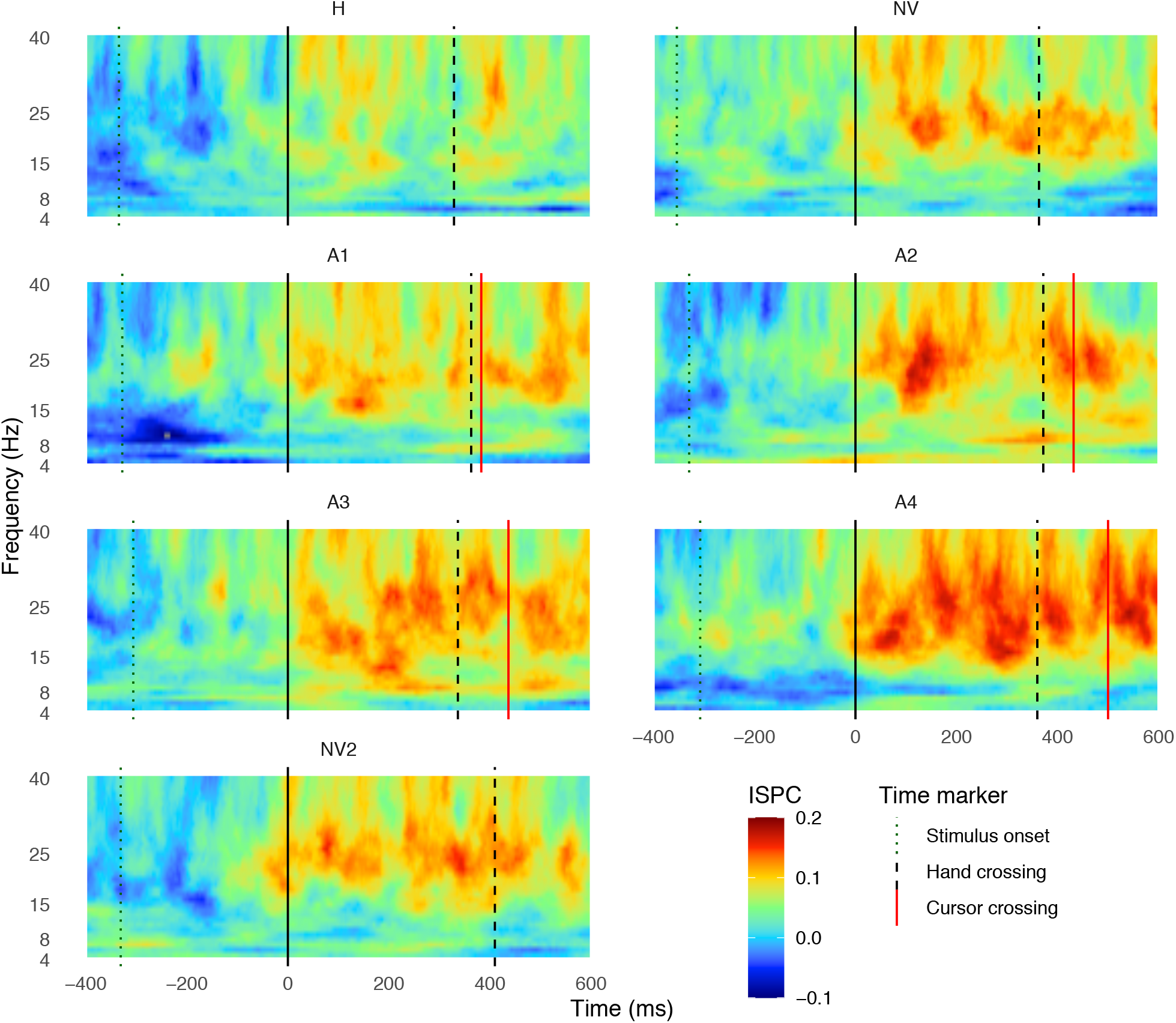
Brain coherence between electrodes in the contralateral motor area (C3) and the visual cortex (O1/2). Heat maps show the time-frequency window of the averaged ISPC values (averaged over trials and participants), with respect to the baseline, between electrodes C3-O1 and C3-O2, as a function of time, for each condition and block (in different panels). The frequency range selected was 4-40 Hz. The time window plotted ranges from 400 ms before the hand reaction time (time zero, solid black line) to 600 ms after this event. The vertical lines indicate the averaged time at which the targets appeared (Stimulus onset - dotted line), the hand RT (continuous black line), the hand movement time (Hand crossing-dashed black line) and the time when the cursor crossed the target line (Cursor crossing - red line).

**Figure 3.**
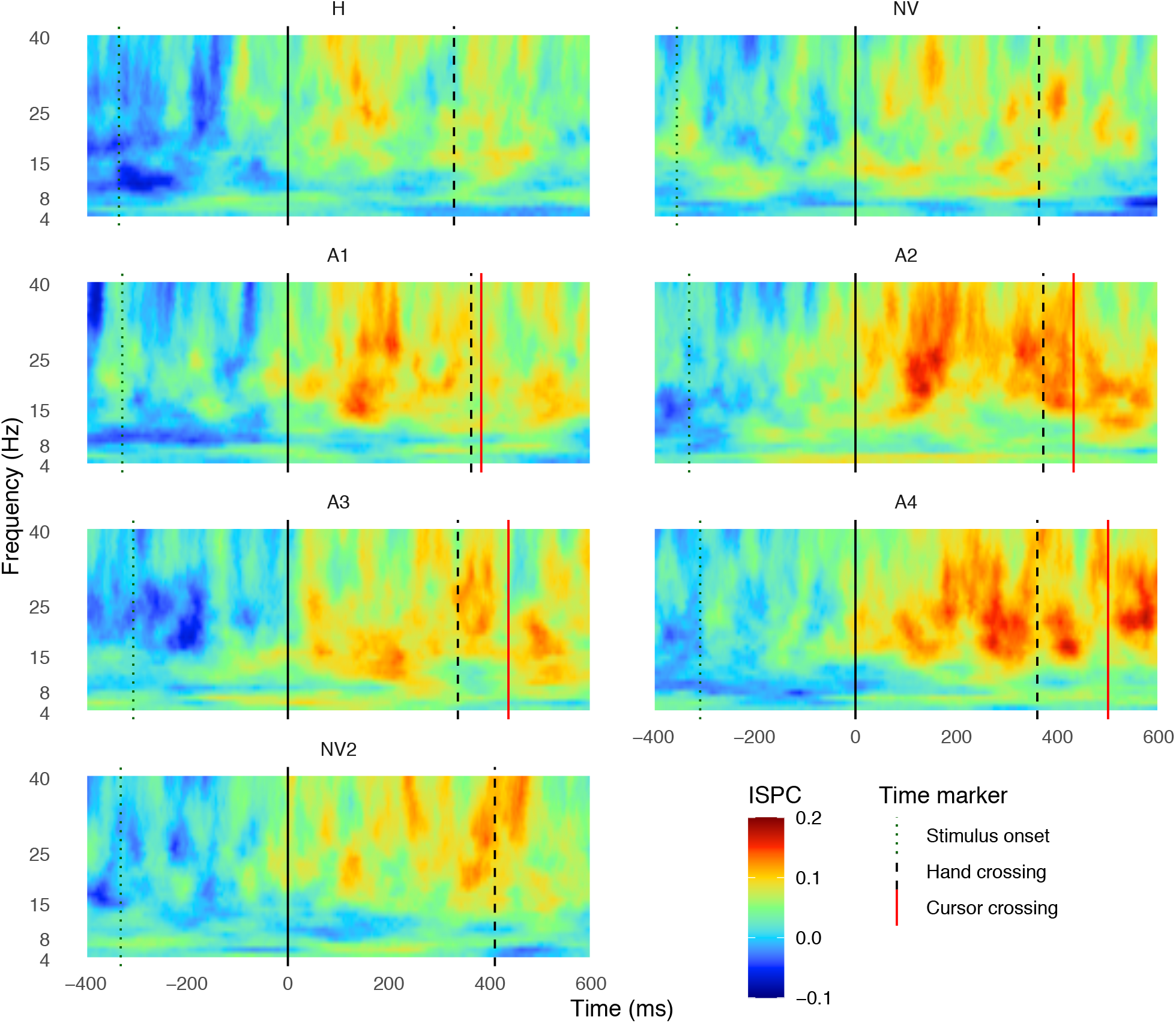
Brain coherence between ipsilateral motor area (C4) and the visual cortex (O1/2). Heat maps show the time-frequency window of the averaged ISPC values (averaged over trials and participants), respect to the baseline, between electrodes C4-O1 and C4-O2, for each condition (in different panels). Details as in Figure 2. The coherence between C4 (ipsilateral motor areas) and O1/2 (visual area) is lower than the one between contralateral motor and visual areas.

Coherence values between C3 and O1/2 (Figure 2), which are associated with the activity of contralateral (left) motor area and visual area, respectively, are low before the hand movement onset (vertical black line, time = 0 ms) in all conditions. In the interval between the stimulus onset and the hand RT, the coherence between both areas increased mainly in beta (16-25 Hz) and gamma (26-40 Hz) bands, which may reflect motor commands setting a proper context for network communication (e.g. increments in beta band activity during tonic muscle contractions and in gamma band due to changes in motor output; see review [40]). Around the time the hand started moving, coherence increases for all conditions. The increase in coherence for the 4 blocks of the A condition appears to be higher than in the other conditions where there is no delayed visual feedback. Furthermore, the longer the delay, the longer the time after interception during which coherence values remain high (e.g. compare A1 and A4). This result is consistent with relevant communication between motor and visual areas in order to control the cursor and coordinate the interceptive movement with the visual target. Similar patterns of coherence were observed between C4 and O1/2 (Figure 3). Importantly though, the coherence pattern in that case (Figure 3) was much weaker than the one between the contralateral motor area and visual area (Figure 2), which indicates that coherence patterns are very likely to be related only to the coupling between visual areas and the contralateral motor area, corresponding to the moving hand.

Since the coherence is phase-locked at time zero (corresponding to the hand RT), the increment in the coherence (observed in Figures 2 and 3) is explained by the evoked responses (within the 100 ms after the phase-lock response) caused by the hand starting move. To study whether the increment in coherence across movement time is actually related to the level of temporal adaptation, we categorized participants as high or low adaptors on the bases of the individual slopes from the linear fits of the temporal error across trials (inset in Figure 1C). Therefore, each participant was assigned to either group based on the comparison of their slope with the median slope (with low adaptors being the ones with slopes smaller than the median slope). Since the individual slopes do not follow a bimodal distribution, we selected the participants with the 6 larger and 6 smallest slopes and grouped these participants in two different groups (labeled as highest and lowest adaptors, respectively) in order to capture the characteristics of low and high adaptation better. We then averaged the coherence activity values for both groups of adaptors for each condition and frequency band (different panels in Figure 4) for the coherence measured at the contralateral motor area (C3; Figure 2) and at the ipsilateral motor area (C4; Figure 3).

**Figure 4.**
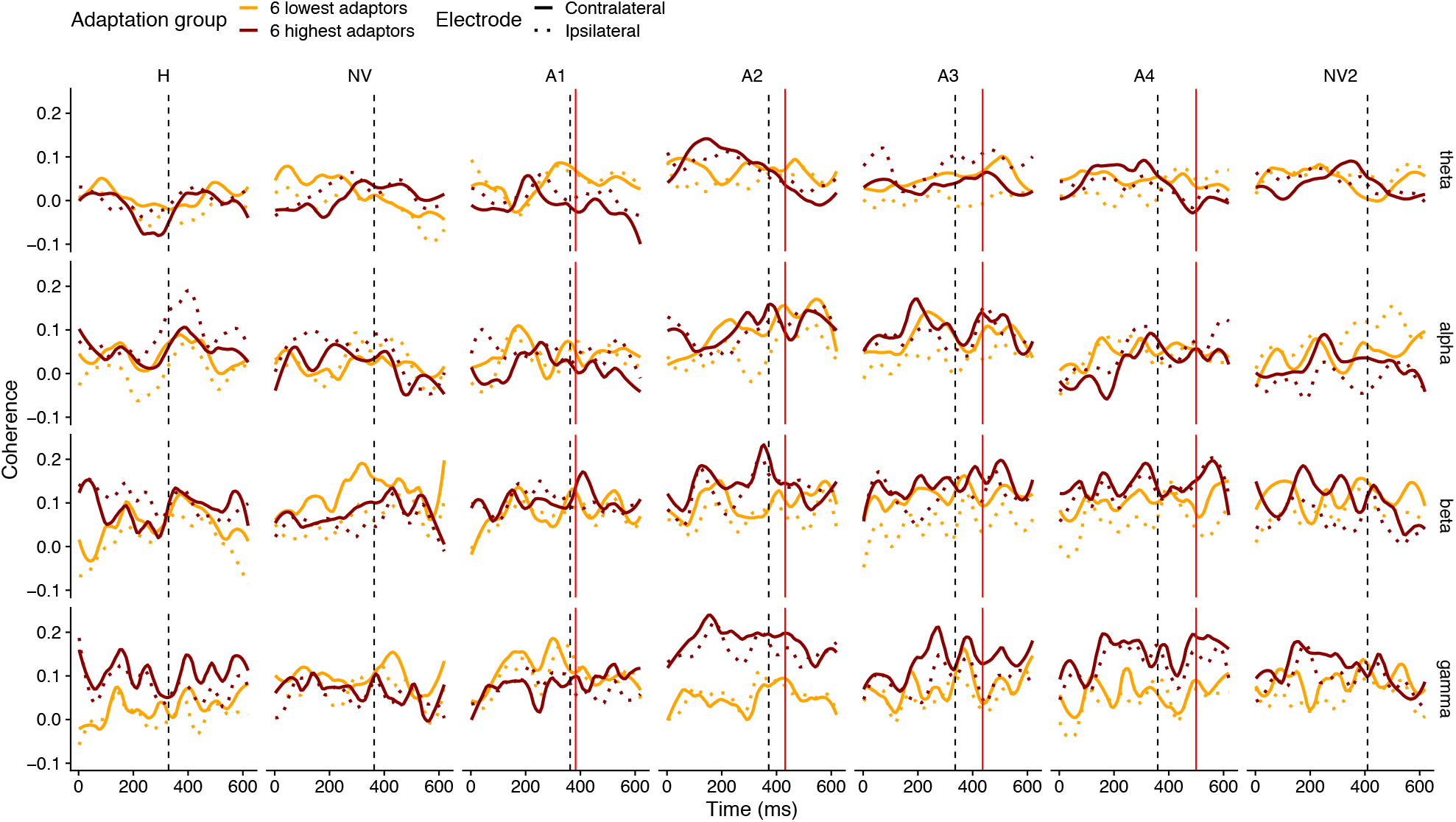
Neural coherence for both groups of participants depending on the level of adaptation. Averaged coherence across the 6 participants with the highest and the 6 participants with the lowest adaptation (red and yellow lines, respectively) as a function of time (0 corresponds to the hand RT), for the different conditions (columns) and frequency bands (rows). The type of line (solid vs dotted) codes the electrode: the coherence between electrode C3 (contralateral motor areas) and the visual cortex (solid line); and between electrode C4 (ipsilateral motor area) and the visual cortex (dashed line). The vertical lines denote the time at which the hand (black dashed) and cursor (red solid in the A condition) crossed the target’s path for the two groups.

By visually inspecting Figure 4, one can see that there are clear differences in coherence between the two groups of adaptors for the high frequency bands (beta and gamma). These differences are even more pronounced during the adaptation blocks, mainly from A2 to A4. In order to determine whether these differences are significant we averaged the coherence (shown in Figure 4) across a time window of 400 ms from hand movement onset for each participant, electrode and condition separately (Figure 5). The length of such temporal window was selected in order to include the relevant time of the interception trajectory [21]. We then conducted an ANOVA on the time-averaged coherence with Adaptation level (lowest vs highest adaptors) as between factor and with site of coherence or electrode (C3: contra-lateral, C4: ipsilateral) and condition (7 levels: from H to NV2 as shown in Figure 5) as within factors. We ran the ANOVA for each frequency band separately. Only beta and gamma band yielded significant main effects or interactions. In the theta band the main effect caused by condition reached marginal significance (F(6,60) = 1.9, p=0.089) reflecting the peak of coherence in condition A2. In the beta band, the site of coherence was significant (F(1,10) = 5.5, p=0.041) due to the larger coherence for the contralateral (C3) electrode (coherence mean value of 0.1 in contrast to the 0.07 obtained for the ipsilateral electrode (C4)). Also, the interaction between site of coherence and condition was significant (F(6,60) = 3.0, p=0.013), while the interaction between adaption level and site of coherence did not reach significance (F(1,10) = 3.3, p=0.099). For gamma band, the site of coherence (contra vs ipsilateral) yielded a significant effect (F(1,10) = 5.4, p=0.043) reflecting the average higher coherence for the contralateral electrode than for the ipsilateral one (coherence mean values of 0.09 and 0.07). Importantly, site of coherence showed a significant interaction with adaptation (F(1,10) =10.4, p=0.009) due to the larger difference between electrodes in the high adaptor group. The fact that both beta and gamma frequency bands show significant differences in coherence caused by the electrode (higher coherence between C3 and O1/2 than the coherence between C4 and O1/2) reflects that coherence is functionally related to the motor task since all the participants performed the task using the right hand. Most importantly regarding adaptation, there was a significant interaction between adaptation level and condition (F(6,60) = 2.6, p=0.025) caused by the increase of coherence in gamma frequency in the adaptation phases (A1 to A4) for high adaptors. This significant effect denotes the relevance of coherence in gamma band for the temporal adaptation.

**Figure 5.**
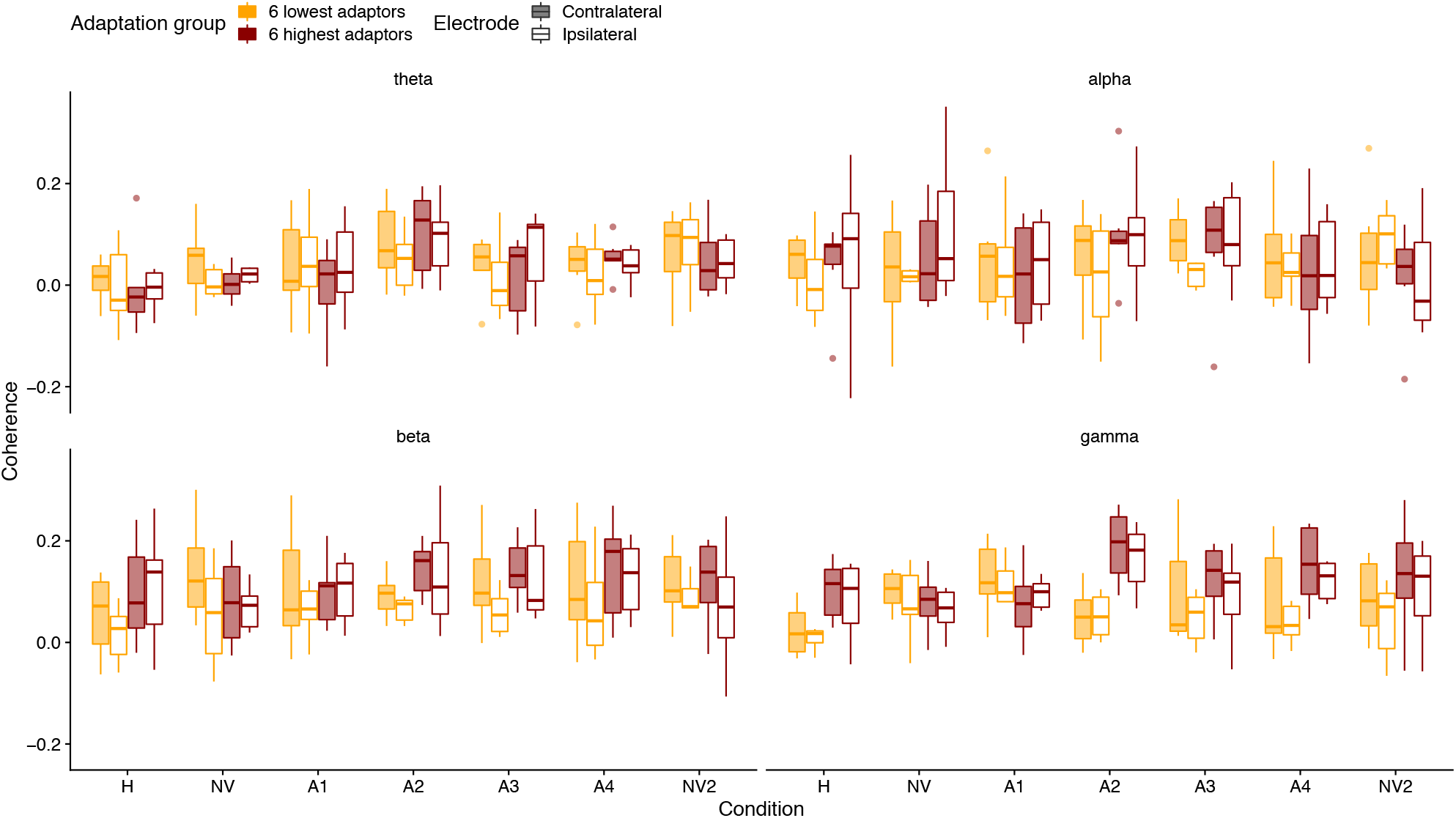
Averaged coherence value across feedback condition. Average coherence for contralateral and ipsilateral motor areas (coded with transparency) over a time window of 400 ms, starting at the hand RT, for high and low adaptors (colour coded) for each feedback condition. Each panel corresponds to a frequency band. In the boxplot, the lower and upper hinges correspond to the first and third quartiles (the 25th and 75th percentiles) and the horizontal line is the 50th percentile. The upper whisker represents the largest value no further than 1.5 · IQR while the lower whisker denotes the smallest value within −1.5 · IQR. Dots represent outliers.

We studied whether the significant interactions reported above were caused by specific conditions exerting an expected effect. We did so by performing planned multiple comparisons corrected for the false discovery rate (Bonferroni correction). Consequently, the p-values reported next are all corrected. We looked at the interaction between site of coherence and condition in the beta band and ran two contrasts. First, we compared condition H versus A for the two electrodes (C3 and C4) and the difference between conditions was only significant, as expected, for the contralateral electrode C3 (diff = 0.039, t = 2.4, p = 0.041), while the difference was not significant for the ipsilateral electrode C4 (diff=0.022, t=1.387, p= 0.336). The second contrast compared NV vs NV2 also for the two electrodes, expecting no differences. Figure 1C and some previous studies [5, 31] showed that participants’ behaviour in the post-adaptation condition goes back to the same level observed in the pre-adaptation condition. As expected, the difference between NV and NV2 was not significant for both sites of coherence: (C3: diff=−0.008, t=−0.403, p=1.0; C4: diff=0.005, t=0.258, p=1.0).

Concerning the expected relevance of the gamma band on temporal adaptation, we saw that the increase in coherence was clearly higher for participants that showed larger values of adaptation. In order to have a closer look at the significant interaction between adaptation level and condition, we contrasted H vs A for each level of adaptation and the difference was significant for the high adaptors (diff= 0.04, t = 2.7, p = 0.018) while it was not for the low adaptors (diff=0.023, t=1.45, p=0.295). Finally, a significant interaction between level of adaptation and site of coherence was also reported for the gamma band. The *post-hoc* analysis revealed that the contralateral (C3) motor area shows a significant difference between the lowest and highest adaptors (diff=−0.083, t = −2.7, p = 0.024), but not in C4 (diff=−0.03604, t= −1.153, p=0.276).

These analyses indicate that only the gamma band drives differences among the participants’ behaviour in terms of the level of adaptation.

### The time course of the simulated coherence

Figure 4 showed the dependence of the level of adaptation to visual delays on the coherence at gamma frequency. Particularly, it shows how gamma coherence increases for high adaptors after movement onset reaching maximum values before the moment of interception (denoted by solid vertical lines). This pattern might mediate an automatic coupling between motor execution and sensory feedback. The fact that the increase in coherence during the movement takes place at gamma frequencies could reflect the need for a closed-loop control to allow possible corrections. Similar automatic coupling for lower frequencies has been reported in [41], where similar rhythmicity appears in motor planning and visual processing areas just before hand movement onset possibly to enhance visuomotor communications. Picking up then the delay between onset of motor actions and the corresponding sensory consequences can be regarded as similar to a detection task whose performance can be enhanced by gamma oscillations [42].

We wonder whether the time course of the increase of coherence could be also described by phase-coupled oscillators. Figure 6A shows the *R* value, reflecting coherence, obtained from the simulations using the Kuramoto oscillators implementing the time delays. The different panels correspond to different simulated delays and the lowest (6 Hz) and largest (40 Hz) simulated frequencies (line type) are shown. The time course of coherence when different numbers of oscillators are simulated are color-coded within each panel. There were no differences between frequencies or delays with regard to the time at which the maximum level of coherence is reached. As it would be expected, the number of oscillators did affect the average time at which the different oscillators were phase-synchronized. Importantly, however, the time at which *R* reaches its maximum value was about 100 ms for 1000 oscillators.

**Figure 6.**
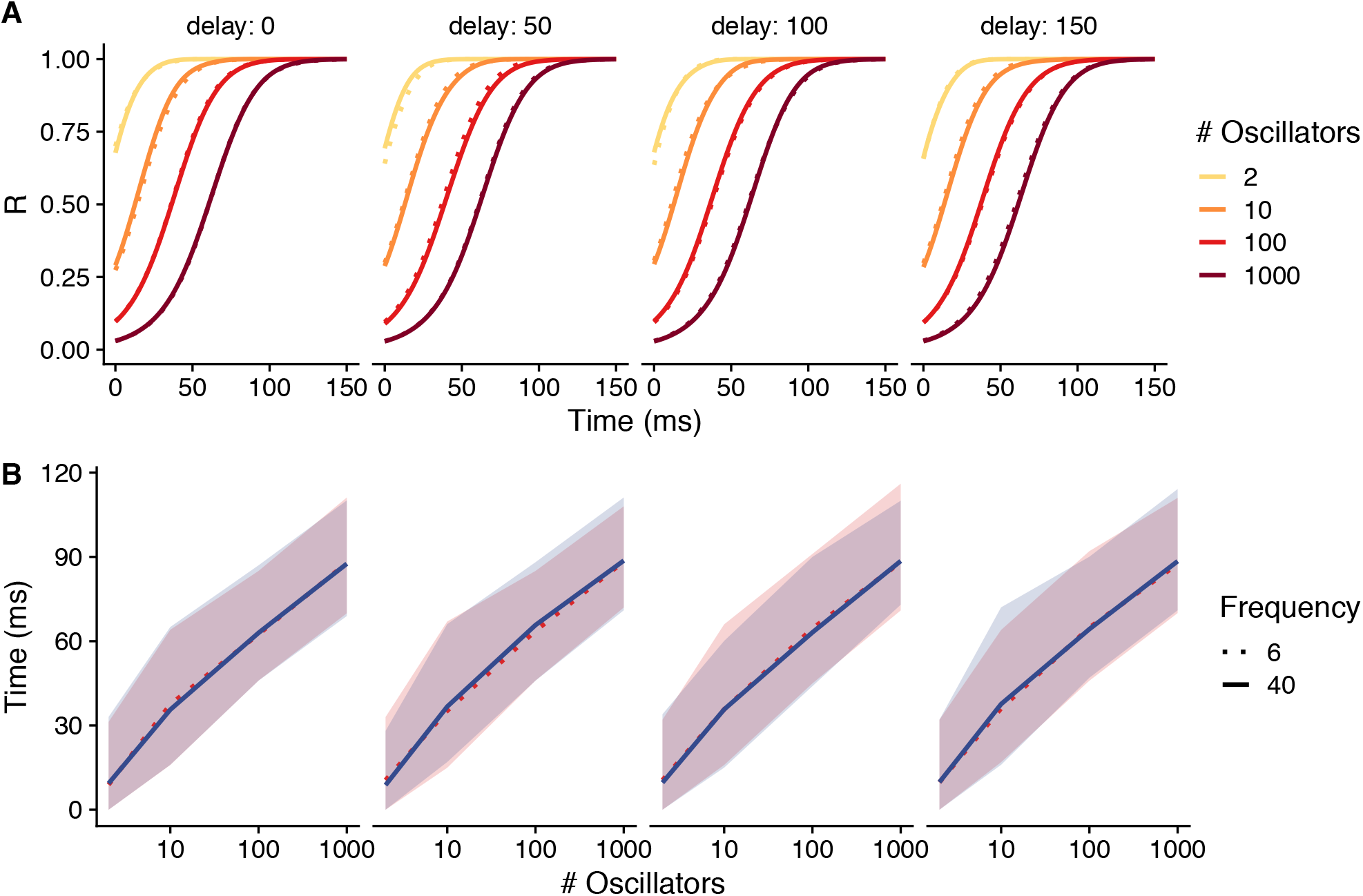
Time course of the Kuramoto oscillator. (A) Temporal dynamics of the phase coupling denoted by the obtained value of phase synchronization (R) which reflects coherence (see Methods for details) as a function of time for simulations involving different number of coupled oscillators (N = {2, 10, 100, 1000}). Different simulated delays are shown in different panels. Since there were no differences between frequencies, we chose to show only 6 Hz and 40 Hz (extreme simulated values) which are coded by different line types. The R values are the means of 1000 simulations. (B) The time at which the maximum phase synchrony is achieved as a function of the different number of simulated oscillators. The value of K was always set to 0.1. The shaded regions denote the 95% CI.

A summary of the time at which the maximum synchrony is achieved as a function of the number of oscillators for each delay and frequencies of 6 Hz and 40Hz can be seen in Figure 6B (note that there were no difference between any frequency bands).

Although we found no difference in the time course of converge among delays (and frequencies), there is clear advantage of communicating at high frequencies if the function is to execute possible correction movements. Figure 7 shows the magnitude of the maximum uncorrected delays as a function of frequency. Obviously, the higher the frequency the smaller the temporal shift due to phase discrepancies. In Figure 7, the grey area denotes the delays that would correspond to the temporal precision due to visual resolution (between 18 and 20 ms). The maximum uncompensated delays in the gamma band (orange regions in Figure 7) already fall within the range of the temporal resolution supported by the visual system resolution. This would further support the functional role of gamma coherence in visuomotor control.

**Figure 7.**
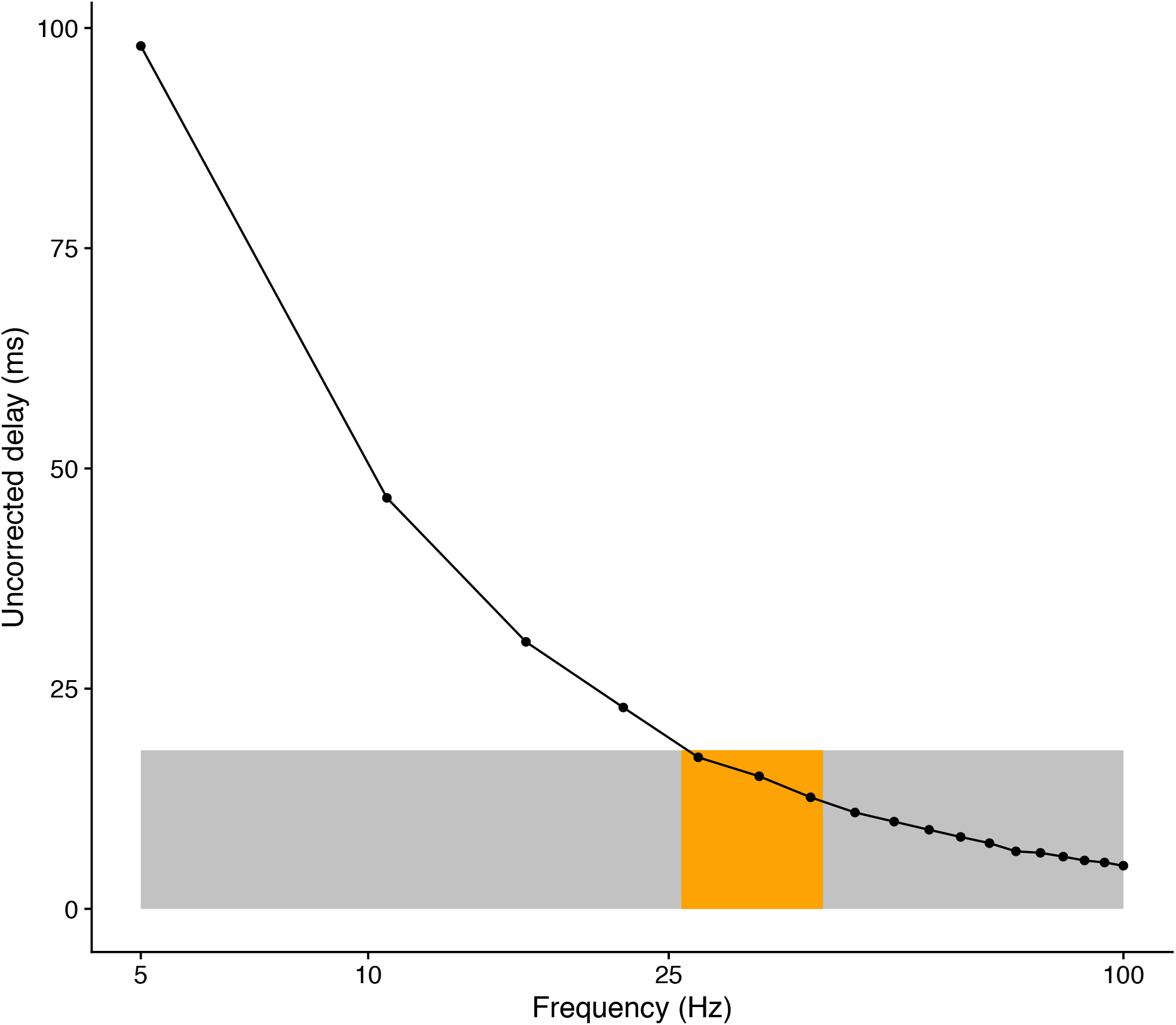
Magnitude of the differential delay that is not corrected for by the Kuramoto oscillator as a function of the oscillation frequency. The grey area denotes the limit of a temporal delay that corresponds to the visual resolution (about 18 ms). The orange region stands for the frequencies in the gamma band.

## DISCUSSION

In this study we recorded electrical brain activity while intercepting a moving target with the hand (visible and not visible) or with a cursor following the hand with a certain delay. Results showed that the neural coherence at high frequencies (particularly at the beta and gamma band) between specific visual and contralateral motor areas was associated with the level of adaptation when participants were exposed to the delayed visual feedback. In addition, those participants who better adapted to external delays presented higher gamma oscillatory synchronization. To our knowledge, this study shows first time evidence that phase synchronization of visual and motor areas in the beta and gamma frequency ranges correlates with the adaptation to visuomotor delays enabling the adjustment of hand movements to successfully intercept a target. Furthermore, we have shown that the time course of this synchronization can be described using a Kuramotor oscillator approach.

The involvement of beta oscillations in the context of motor experiments is expected, as several studies have reported beta activity in the preparation and initiation of movements [11, 43] as well as after movements signaling the magnitude of the error [13] or the uncertainty associated with the forward model of the movement [14]. It is important to note that the majority of previous studies have focused on the power of the beta oscillatory activity, that is, the increase of energy at this frequency band. In the present paper we show that this oscillatory activity plays a key role in the coordination of some of the areas responsible of the visuomotor temporal adaptation. Interestingly, not only beta, but also gamma activity is responsible of such synchronization. The pattern of adaptation shown in Figure 1C indicates that people detect the progressively increasing discrepancy between the hand movement and the visual feedback. In this sense the adaptation would likely require a conflict monitoring mechanism consistent with activation of frontal areas (e.g. anterior cingulate cortex; see [44]). This activation has been observed in temporal recalibration [1] when the timing of the visual feedback after a key press was perturbed. In the case of a key press, adaptation could be based on trial-based corrections of the previous error [45] in a similar way as we learn to adjust continuous movements towards spatially distorted targets [12, 13]. These previous studies have identified the power of beta oscillations after the end of a movement as a signature of the magnitude of the error (i.e. lower beta power for larger errors). This suggests an important role of beta oscillations in trial-to-trial learning from errors in consolidating an internal model. Our participants, regardless of the level of adaptation, do not show a change of coherence (nor power) in the beta band during the adaptation phase (see Figure 4), although the coherence in this band was larger for those who adapted more. However, it is not clear that we can associate the smaller beta activity to larger errors at the end of the trial that are induced by our temporal perturbation because our perturbation was incremental. Thus, with this kind of perturbation it might take some trials for people to detect a temporal discrepancy between the timing of the expected feedback and the actual visual feedback. A recent study by [14] also reports a negative correlation between beta activity and the uncertainty associated with a forward model estimate. This is relevant for our study because it considers the history of previous errors. On the other hand, computational Bayesian models of error-based learning (e.g.: [38, 45, 46]) predict that higher uncertainty of the internal model estimate would lead to larger learning rates. In part, the smaller beta activity in the coherence (and supposedly larger model uncertainty) in all conditions for participants that adapt less in our task would be consistent with larger final errors that arise when one does not adapt (i.e. the need to update an internal model).

The adaptation mechanisms involved in the visuomotor activity have to be fast by nature as bi-directional coherence between sensory and motor areas, irrespective of the putative communication mechanism, must enable the ability to make fast sensorimotor correction when a discrepancy in the sensory inflow with the prediction is detected. The mechanism supporting this process could share principles with the improvement of change detection of visual stimuli due to gamma coherence between V1 and V4 [47]. Humans show high capacities for fast online corrections in reaching and interception based on visual feedback [48]. The involvement of beta and gamma bands in timing visuomotor actions appears then to be meaningful in the context of our sensorimotor task but at the same time challenges some models on brain connectivity. Some theoretical accounts propose that the synchronization of distant areas is better driven by low oscillatory activity (such as alpha and low beta bands) and that high frequency bands would be more appropriate for short-range (more local) network communication [49, 50]. The reason for that is that the connectivity depends on the conduction velocity between the connected areas and the synchronization of distant areas might be done either by using fast connection or decreasing the oscillatory frequency [51]. However, these low frequency bands do not seem to be appropriate for the fast corrections required in visuomotor synchronization. Importantly, simulations using Kuramoto oscillators show that the time course of phase synchronization was not different for the different frequency bands that we simulated. However, we used a type of oscillator that approximates the temporal delay between oscillators. This approximation can successfully cope with delays only at high-frequency oscillatory activities (see Figure 7) because the magnitude of delay that is not accounted for is negligible for high frequencies. Therefore, with the inclusion of a mechanism dealing with delays by means of coupled oscillators, high-frequency oscillatory activity emerges as clear possibility for the fast coordination of distant areas needed to adapt to external delays. Importantly, previous experiments using other tasks have also showed that the coherence at high frequency bands is able to coordinate distant cortico-cortical distances (see for example [52–54]).

It is important to note that our results do not allow to easily disentangle the role of gamma and beta activities in the present experimental design. One possible differential role of the gamma band activity is suggested by the modelling of the Kuramoto oscillator. At the time of hand RT a trigger would start a process of oscillatory phase coupling that would end up in two different population of neurons firing in sync taking into account the traveling delays. Gamma coherence would then support efficient communication between sensory feedback and motor areas within the trial. Figure 4 shows peaks of gamma coherence as early as about 200 ms after hand onset for high adaptors.

Participants that adapt better to the delays move for a longer time (about 377 ms) than those that adapt less (about 313 ms) in the last adaptation phase. Considering the time at which maximum coherence is achieved in this phase (about 200 ms), participants that adapt more have a more comfortable time window (about 177 ms versus 113 ms) to control the delayed cursor and make within trial corrections before it is too late. The interception task used in this study had high spatio-temporal constraints and would certainly benefit from bi-directional fast communication between visual and motor areas within a single trial. In addition, the frequency at which the gamma band operates would allow fast monitoring of the visual feedback consistent with the temporal resolution of the visual system [48]. The capability of fast corrections despite the built-in delays in sensorimotor loops is often attributed to the role of forward models with the ability to integrate sensory inflow and motor output [55]. Therefore, one can regard the ability to adapt to additional delays as extending the predictions of forward models.

One issue to be cautious about is whether the network communication in the adaptation to delays is one-way (i.e. from motor areas to visual areas) so that the sensory inflow is timely processed or bi-directional. The communication through coherence (CTC) hypothesis is still a matter of debate. Recent proposals [56] invoke two unidirectional processes with non-zero phase synchronization rather than bi-directional communications with zero-phase [23, 57]. Without aiming at providing a computational model of the temporal adaptation process, the reported simulations in this study do provide some insights into the regulation mechanisms to compensate for phase delays. We want to point out that a simple implementation of a phase correction mechanism results in a clear advantage for high frequencies (Figure 7). We can assume that this mechanism is implemented across larger networks and hence many more oscillators in the brain might increase the time that is needed to couple their phases. When time is in short supply in sensorimotor tasks with moving objects like ours, the temporal requirements for phase coupling could have rendered this mechanism implausible. However, our simulations with a relatively large number of oscillators (1000) indicate that the mean time needed to reach a maximum coherence level is still within very reasonable values (about 100 ms) (Figure 6B) independently of the delays. The Kuramoto model with spatial embedding that we have used would still support performance in sensorimotor tasks requiring a precision of 12 ms (Figure 7).

## CONCLUSION

We have shown that an increase of coherence in the gamma band during the movement predicts the level of adaptation to temporal delays. We interpret this result as the gamma band supporting fast communication between motor and sensory areas enabling fast corrections when controlling the delayed cursor.

## Acknowledgements

This work was supported by project PSI2017-83493-R (AEI/FEDER, UE). The first author was supported by fellowship BES −2014 −069289. JMP is partially supported by the ICREA Academia grant. The authors thank Matthias S Keil for his advice in the computer simulations.

## Authors Contributions

JLM designed research; CC performed research; JLM, JMP, CM and CC analysed data; JLM, JMP, CM and CC wrote the paper; and CC, CM, JMP and JLM approved the paper.

## Data availability

Raw data can be found in https://osf.io/6jbqx/.

